# Transformations in prefrontal ensemble activity underlying rapid threat avoidance learning

**DOI:** 10.1101/2024.08.28.610165

**Authors:** Christopher J. Gabriel, Tanya Gupta, Asai Sanchez-Fuentes, Zachary Zeidler, Scott A. Wilke, Laura A. DeNardo

**Affiliations:** Department of Physiology, University of California, Los Angeles, Los Angeles, United States; UCLA Neuroscience Interdepartmental Program, University of California, Los Angeles, Los Angeles, United States; Department of Psychiatry, University of California, Los Angeles, Los Angeles, United States

## Abstract

The capacity to learn cues that predict aversive outcomes, and understand how to avoid those outcomes, is critical for adaptive behavior. Naturalistic avoidance often means accessing a safe location, but whether a location is safe depends on the nature of the impending threat. These relationships must be rapidly learned if animals are to survive. The prelimbic subregion (PL) of the medial prefrontal cortex (mPFC) integrates learned associations to influence these threat avoidance strategies. Prior work has focused on the role of PL activity in avoidance behaviors that are fully established, leaving the prefrontal mechanisms that drive rapid avoidance learning poorly understood. To determine when and how these learning-related changes emerge, we recorded PL neural activity using miniscope calcium imaging as mice rapidly learned to avoid a threatening cue by accessing a safe location. Over the course of learning, we observed enhanced modulation of PL activity representing intersections of a threatening cue with safe or risky locations and movements between them. We observed rapid changes in PL population dynamics that preceded changes observable in the encoding of individual neurons. Successful avoidance could be predicted from cue-related population dynamics during early learning. Population dynamics during specific epochs of the conditioned tone period correlated with the modeled learning rates of individual animals. In contrast, changes in single-neuron encoding occurred later, once an avoidance strategy had stabilized. Together, our findings reveal the sequence of PL changes that characterize rapid threat avoidance learning.

## INTRODUCTION

The capacity to avoid threats, particularly those with dire consequences, is especially critical for survival. Individuals that learn slowly are unlikely to survive very long. Thus, animals must rapidly learn to avoid aversive outcomes by predicting threats and taking preemptive actions to avoid them. Often, this means identifying locations that are safe in the context of specific, impending threats and then remaining in those locations until the threat has passed. Thus, animals quickly learn how threat-predicting cues alter the implications of entering or leaving a safe location. Most studies to date have focused on the neural underpinnings of well-established behavioral strategies for avoiding threats. But the neural computations underlying rapid and flexible threat avoidance learning remain poorly understood.

The medial prefrontal cortex (mPFC) integrates information about emotionally salient cues to promote adaptive behavioral responses (Euston et al., 2012; Giustino & Maren, 2015; Mack et al., 2024). On the other hand, mPFC dysfunction is often involved in mood and anxiety disorders characterized by excessive avoidance of perceived threats (Clauss et al., 2016; Mack et al., 2023; Xu et al., 2019). Activity in the prelimbic subregion of mPFC (PL) encodes threatening cues and threat-induced behaviors, including freezing and avoidance behaviors (Burgos-Robles et al., 2009; Courtin et al., 2014; Cummings & Clem, 2020; Diehl et al., 2018; Jiao et al., 2015; Kajs et al., 2022; Moscarello & LeDoux, 2013; Zelikowsky et al., 2014). In well trained mice, PL activity is required to actively avoid a signaled threat (Bravo-Rivera et al., 2014; Diehl et al., 2018, 2020) and PL activity accurately predicts whether mice will successfully avoid aversive outcomes (Ehret et al., 2024; Jercog et al., 2021). While these studies show that PL is a key player in threat avoidance, they largely focused on well-established behavioral strategies. Thus, how the necessary neural dynamics arise during learning – especially when learning occurs rapidly, in just a few trials – is poorly understood.

To better understand the processes underlying this rapid learning, we used head-mounted miniscopes (Cai et al., 2016; Ghosh et al., 2011) to record Ca^2+^ signals from PL as mice learned platform mediated avoidance (PMA). In PMA, a conditioned tone prompts mice to navigate to a safety platform to avoid a footshock (Bravo-Rivera et al., 2014; Diehl et al., 2019). We compared PL activity patterns throughout PMA learning with PL activity in mice exploring the same context, but without shocks. Over the course of learning, we identified a rapid shift in the PL population code that preceded changes observable in single-neuron encoding. Early in the learning process, PL population dynamics accurately predicted trial outcomes and precisely tracked individual learning rates. Once behavioral performance stabilized, neurons that encoded avoidance behaviors or risky exploration were strongly modulated by the conditioned tone, and neurons encoding the onset of the conditioned tone were strongly modulated by the location of the mouse (on or off the safety platform). Together our findings reveal that during avoidance learning, PL rapidly generates novel representations of whether mice will take avoidance or exploratory actions during an impending threat. Moreover, we reveal the sequence of changes that unfold in PL and how they relate to individual learning rates.

## RESULTS

### Tracking changes in PL neuronal activity during signaled avoidance learning

To examine the neural correlates of avoidance learning, we used miniscopes to record Ca^2+^ activity from hundreds of PL neurons as animals learned PMA (Figure 1A,B and S1). All mice were trained for 2 days in PMA (Figure 1D). On day 1, the first 3 tones were presented without shocks to assess baseline tone responses from the recorded cells. The remaining 9 tones co-terminated with a mild 2-second foot shock. On day 2, all 12 tones were paired with shock. To determine if the observed changes in neural dynamics were due to avoidance learning, we compared with PL activity recorded from non-shocked (NS) control mice that underwent the same procedures, but without receiving any foot shocks. We imaged PL activity during PMA on days 1 and 2 and extracted calcium signals associated with individual neurons using a constrained non-negative matrix factorization algorithm (Dong et al., 2022). We then analyzed single neuron and population level activity patterns. We recorded a total of 1394 PL neurons across 9 shocked animals, and 967 PL neurons across 7 NS animals.

**Figure 1.**
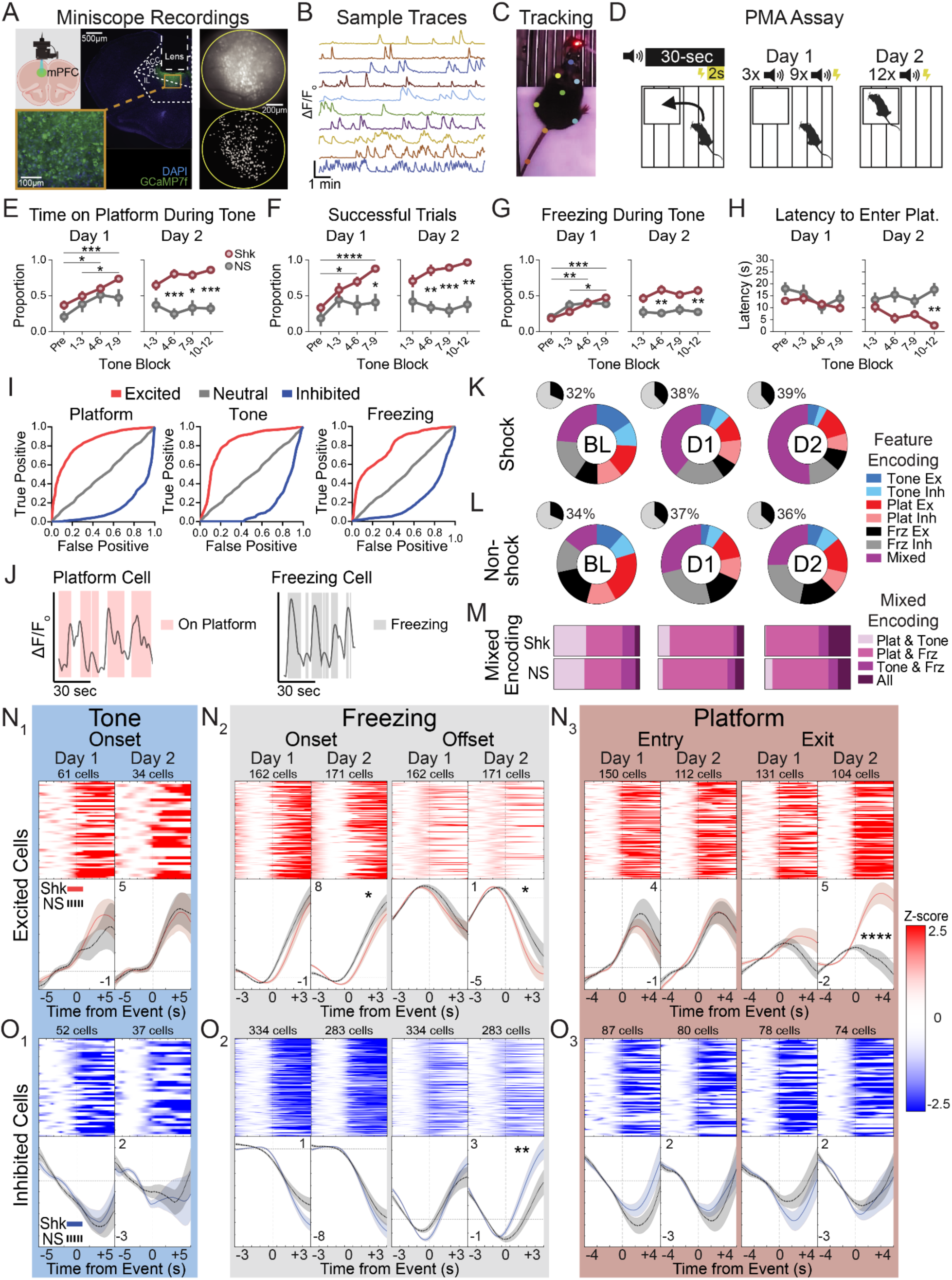
Changes in PL Encoding as Animals Learn PMA. A. Schematic showing virus and miniscope imaging targeting mPFC (image generated by BioRender), GCaMP7f expression, GRIN lens placement, maximum intensity projection of example miniscope window showing recorded neurons *(top right)* and extracted cell spatial footprints *(bottom right)*. B. Example neural traces recorded from PL, displayed in ΔF/Fo. Traces show 10 minutes of recording. C. Example of DeepLabCut-based pose estimation on a mouse wearing a UCLA miniscope. D. PMA assay design. (E-H) Summary of PMA behavior data on Day 1 (left) and Day 2 (right) for shocked and non-shocked groups. Asterisks above plots indicate significant differences between Shock tone blocks. Asterisks between groups indicate significant differences between Shock and NS groups. Comparisons not shown were not significant. (Two-way ANOVA; Day 1: Shk: N=11, NS: N=9; Day 2: Shk: N=8, NS: N=8). E. Time on platform during tone. F. Successful trials. G. Freezing during tone. H. Latency to enter platform. I. Example ROC-based excited, suppressed, and neutral cells for platform, tone, and freezing. J. Example traces from significantly responsive cells overlaid with behavior labels. K. Summary of significantly responsive neurons in shocked animals across PMA. Black/gray pie charts show percent of recorded cells that were active. Multi-colored pie charts show profiles of tone, platform, and freezing cells. (Two-sided chi-square with Yates’ correction for mixed cells on day 2; Chi: 43.13, df: 1, z: 6.567, p<0.0001). Time epochs: BL (day 1, tones 1-3); Day 1 (tones 9-12); and Day 2 (tones 9-12). Cells analyzed: BL/Day 1: 1394, Day 2: 1031. L. Same as K for non-shocked animals. Cells analyzed: BL/D1: 847, Day 2: 967. M. Bar plots of mixed response cells shown in K & L. (N,O) *(Top)* Summary of cells that were significantly responsive during tone, freezing, and platform transitions, separated by excited and inhibited responses. Heatmaps show cells pooled across shocked animals. *(Bottom)* Mean activity of significantly responsive cells pooled across groups and separated by response type (excited or inhibited; asterisks indicate significance on Welch’s t-test of pooled AUC values). N. Heatmaps and mean response plots for significantly excited tone, freezing, or platform cells. O. Same as N for inhibited cells. *P<0.05, **P<0.01, ***P<0.001, ****P<0.0001. Graphs show mean ± SEM. See Supplemental Table 1 for additional statistical information.

To precisely quantify learning-related behavior, we tracked mice using automated approaches and extracted detailed behavioral metrics (Figure 1C; Gabriel et al., 2022; Mathis et al., 2018). On day 1, shocked mice exhibited a steady increase in the time spent on the platform, in the proportion of successful trials, and in freezing levels during the tone (Figure 1E-G). We defined successful trials as those when mice accessed the platform before the beginning of the shock and remained on the platform for the duration of the shock period (i.e., until the end of the tone). By the end of day 1 and for all of day 2, shocked mice had significantly more successful trials compared to non-shocked controls (Figure 1F). On day 2, shocked mice also spent significantly more time on the platform and more time freezing during the tone compared to non-shocked controls (Figure 1E,G). Mice in both groups had a comparable latency to enter the platform after tone onset on day 1, but shocked mice had a significantly shorter latency to enter by the end of day 2 (Figure 1H).

### Avoidance learning drives an increase in the proportion of PL neurons with mixed selectivity

Studies of aversive learning have identified changes in single neuron encoding in PL, including more neurons that encode conditioned tones, neurons whose activity correlates with sustained freezing behavior, neurons that suppress their firing during avoidance actions, and more (Burgos-Robles et al., 2009; Diehl et al., 2018). However, these studies primarily described neural activity during well-instantiated signaled avoidance. To determine how and when such changes emerge remains poorly understood, we examined single-neuron encoding of PMA features on day 1 of PMA, early in learning, versus on day 2, when behavioral performance becomes more stable. We first investigated whether and when individual PL neurons are significantly tuned to features of PMA and how their encoding changes with learning. To do this, we use receiver operating characteristic (ROC) analysis (Kingsbury et al., 2019; Li et al., 2017) to determine whether individual PL neurons were reliably excited or inhibited in response to each feature (Figure 1I,J). We computed cell tuning properties during three periods: the baseline period of day 1 (BL; tones 1-3), the end of day 1 (D1; tones 9-12), and the end of day 2 (D2; tones 9-12). We examined features in three categories: tone (including responses to both tone onset and tone offset), platform (including responses to platform entries, exits, and location), and freezing behavior.

In each period (BL, D1, and D2), the overall fraction of PL cells that were significantly responsive to PMA features was similar between groups. In shocked animals, we observed a modest increase from 32–39% significantly responsive PL neurons (black/gray pie charts, Figure 1K). Non-shocked control mice exhibited a similar trend (34–36%, black/gray pie charts, Figure 1L). On day 1, shocked and non-shocked control mice had similar proportions of PL neurons that were tuned to the tone, the safety platform, freezing behavior, or their combinations (colored wheel plots, Figure 2K,L). In contrast, by day 2 of PMA, shocked mice had a significantly higher proportion of PL neurons that exhibited mixed selectivity (i.e., that were significantly modulated by multiple features of PMA; shocked: 51%, non-shocked: 25%; Χ^2^: 43.13, p<0.0001) (colored wheel plots, Figure 2K). While we observed different combinations of mixed selective tuning (e.g., tone+platform cells, tone+freezing cells, etc.), shocked mice exhibited more PL neurons that encoded the combination of the tone, platform, *and* freezing behavior (Figure 1M and S3).

**Figure 2.**
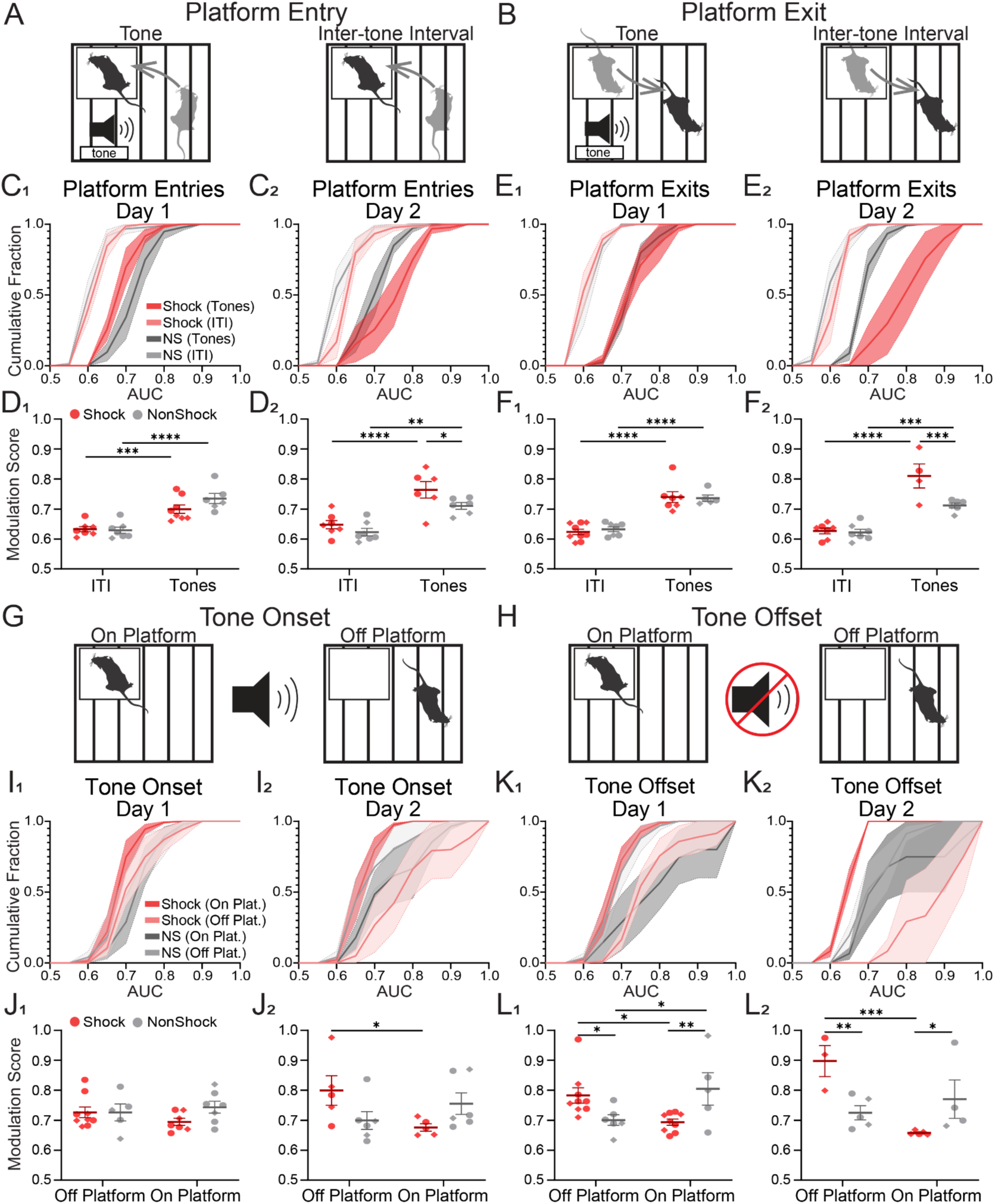
Enhanced PL Modulation During Tone and Platform Activity. A. Depiction of platform entries during tone and inter-tone intervals (ITIs). B. Depiction of platform exits during tone and ITIs. (C-F) Activity levels of behaviorally-responsive cells during tone and ITI. C. Cumulative frequency histograms of auROC values for platform entry responses during tone or ITI. D. Modulation scores during platform entries (Two-way ANOVA; Day 1: Shk: N=9; NS: N=7; Day 2: Shk: N=7, NS: N=7). E. Same as C for platform exit responses during tone or ITI. F. Modulation scores during platform exits (Two-way ANOVA; Day 1: Shk: N=9; NS: N=7; Day 2: Shk: N=6, NS: N=7). G. Depiction of tone onset when animal is on or off platform. H. Depiction of tone offset when animal is on or off platform. (I-L) Activity levels of behaviorally-responsive cells when animals are on or off platform. K. Cumulative frequency histograms of auROC values for tone onset responses while on/off the platform. M. Modulation scores during tone onsets (Two-way ANOVA; Day 1: Shk: N=8; NS: N=7; Day 2: Shk: N=7, NS: N=7). N. Same as I for tone offset responses while on/off the platform. O. Modulation scores during tone offsets (Two-way ANOVA; Day 1: Shk: N=9; NS: N=7; Day 2: Shk: N=5, NS: N=6). *P<0.05, **P<0.01, ***P<0.001, ****P<0.0001. Circles represent female subjects and diamonds represent male subjects. Graphs show mean ± SEM. See Supplemental Table 1 for additional statistical information.

Having identified the PL neurons that encode key features of PMA, we next investigated how the overall activity of these feature-tuned neurons changes across learning. When we calculated the mean activity levels for all cells in each category, we observed no significant differences in the amplitude of excited or inhibited cell responses to tone onset or offset (Figure 1N_1_,O_1_).

However, we did observe significant differences between shock and non-shocked controls for freezing and platform tuned cells. On day 2, shocked mice had a significant decrease in excitation levels following freezing onset and offset (Figure 1N_2_), as well as a significant decrease in inhibition levels following freezing offset (Figure 1O_2_). While we observed no differences in activity during platform entries, shocked mice exhibited a large increase in activity following platform exits on day 2 (Figure 1N_3_,O_3_). Together, these data suggest that while learning does not significantly alter the proportion of PL neurons selectively tuned to specific features, it does impact the overall activity levels of those neurons and selectively enhances the proportion of mixed-selective neurons.

### Avoidance learning enhances the encoding of avoidance actions and safe locations during threatening cues

During PMA, mice can access a safe location both in the presence or absence of an impending threat (tone vs. ITI). This allowed us to study how PL activity encoded safety seeking vs. exploration with respect to threat imminence, and how these activity patterns emerged during learning. Based on our findings of more mixed selective neurons in PL, we hypothesized that through learning, threatening cues begin to alter the encoding of avoidance-related behaviors. To determine if and when such changes occur, we examined PL neurons tuned to platform entries or exits, comparing their activity levels during the tone vs. during the inter-tone interval (ITI) (Figure 2A,B). We quantified activity using area under the curve (AUC) values from ROC plots for both excited and inhibited PL neurons, where cells with larger absolute AUC values are more strongly modulated. We then plotted AUC values as cumulative distributions (Figure 2C,E,I,K). On day 1, neurons encoding platform entries or exits were more strongly modulated during the tone for both shocked and NS control mice (Figure 2C_1_,E_1_). On day 2, however, we observed a further increase in modulation during the tone in shocked mice beyond the level of non-shocked controls (Figure 2C_2_,E_2_). To ensure these patterns were not driven by outlier mice, we also calculated a modulation score for each animal using individual AUC values from the cumulative frequency histograms (see methods; Figure 2D,F). The patterns we observed were consistent across mice: shocked mice had significantly higher modulation scores for both platform entries and exits that occurred during the tones on day 2 (Figure 2D_2_,F_2_).

In a complementary set of analyses, we examined to what extent tone-related activity was modulated based on the animal’s location (ON vs. OFF platform), examining both tone-onset and offset responses (Figure 2G,H). Shocked mice exhibited no location-dependent modulation of the tone-onset responses on day 1, but by day 2, tone-onset responses were more strongly modulated when the animal was OFF the platform (i.e., when threat levels were high; Figure 2I_2_,J_2_). Tone-offset responses were also more strongly modulated when animals were OFF the platform (i.e., had just experienced a foot shock; Figure 2L_1_,L_2_). Non-shocked control mice did not exhibit location-dependent modulation of tone-evoked activity on either day. These data indicate that once animals have learned to avoid a signaled threat, the presence of the threatening cue strongly modulates the encoding of entries and exits from a safe location.

### Population dynamics prospectively represent trial outcomes early in learning

Emergent population dynamics in PL can represent features of behavior that are not apparent at the single neuron level (Powell & Redish, 2014; Siniscalchi et al., 2019). In well trained mice, PL population activity predicts whether animals will successfully avoid a footshock and encodes tone-induced avoidance actions (Ehret et al., 2024; Jercog et al., 2021) and is highly sensitive to threat levels (Casanova et al., 2024). But when these changes emerge during learning is not well understood.

Thus, we investigated whether PL population dynamics could predict trial outcomes during avoidance learning and if so, which neurons contributed to the predictive power. Focusing on early learning (Day 1), we trained support vector machines (SVMs; Awad & Khanna, 2015) to decode the trial outcome from PL activity for each animal. We then evaluated decoder performance continuously across tone-related epochs. Decoder performance using all recorded neurons was generally greater for shocked mice than non-shocked controls, reaching significance for the pre-shock (23-28s after tone onset) and shock (27-32s after tone onset) epochs (Figure 3A). Notably, decoding performance was already better even before the onset of the tone. This suggests that PL activity rapidly forms a representation of the threatening context or fearful state that can predict successful avoidance even during the ITI.

**Figure 3.**
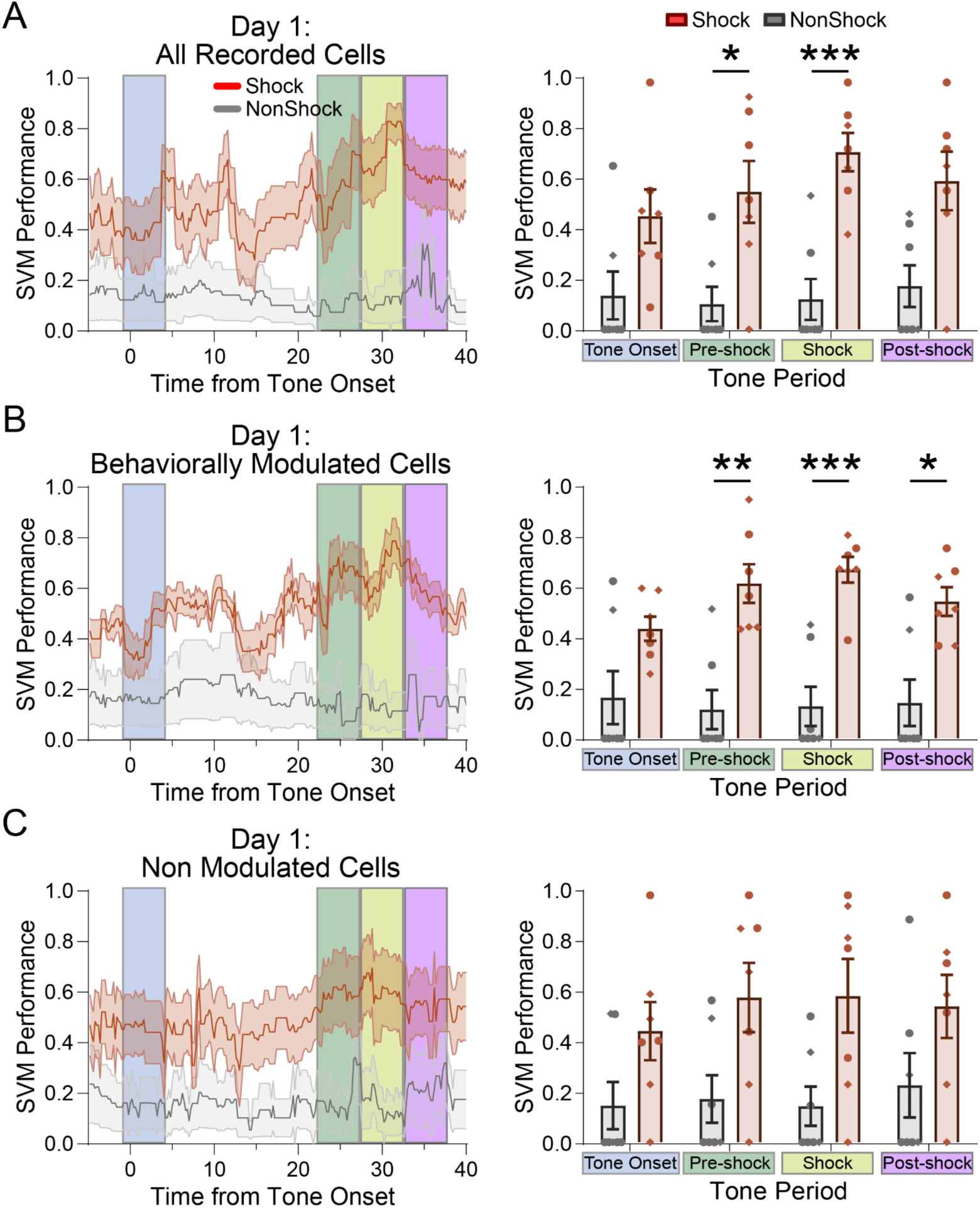
Population Decoding Emerges on Day 1 of PMA. A. (*Left*) Average performance for decoding of trial outcome across the tone for shocked and non-shocked mice. (*Right*) Performance quantification for various tone epochs (Two-way ANOVA; Shk: N=7, NS: N=7). B. Same as A using only cells identified by ROC as behaviorally-modulated (platform, tone, or freezing). C. Same as A using only cells identified by ROC as non-responsive to behavioral features. *P<0.05, **P<0.01, ***P<0.001. Circles represent female subjects and diamonds represent male subjects. Graphs show mean ± SEM. See Supplemental Table 1 for additional statistical information.

We next investigated whether task modulated cells (ROC analysis; Figure 1) contributed more to decoder performance than cells that did not encode features of PMA. Decoder performance in shocked mice was significantly better than in non-shocked controls for the pre-shock, shock, and post-shock epochs (Figure 3B). In contrast, performance was not significantly better in shocked mice when decoding was performed using only the non-modulated PL cells (Figure 3C). So, although changes in single neuron encoding properties were not evident until day 2 of PMA (Figure 1), meaningful changes in their population activity emerged on day 1 (Figure 3). These data suggest that the population dynamics of behaviorally modulated neurons may be a key driver of early learning.

### PL population activity states are related to individual learning rates

We observed some individual variation in decoder performance, which we speculated could be because individual mice learned at different rates. To better understand how PL dynamics relates to PMA learning, we applied a Rescorla-Wagner (RW) learning model to individual behavior and computed the learning rate for each mouse (Rescorla & Wagner, 1972). To determine how mice learn to avoid, we tested three separate models that assume mice learn on trials where they fail to avoid and receive shocks (shock), on trials where they succeed in avoiding a predicted shock (avoid), or on both trial types (hybrid) (Figure 4A). PMA learning was best modeled by the hybrid approach, suggesting that mice learn from both failed and successful avoidance trials (Figure 4B). In the hybrid model, fraction of time ON the platform closely matched observed behavioral data (Figure 4C).

**Figure 4.**
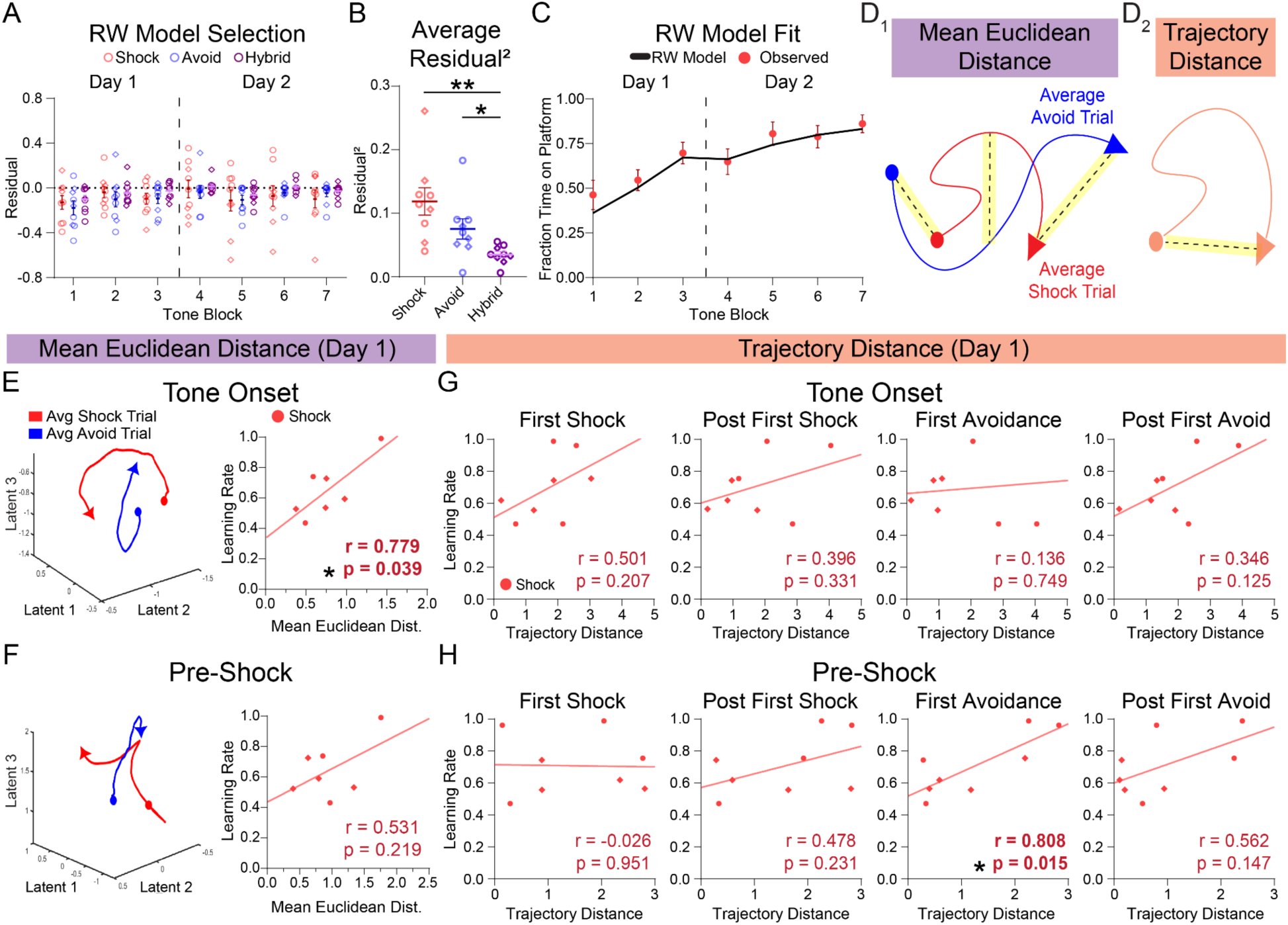
PMA Learning Rate Correlates with PL Population Responses. A. Comparison of residuals between average predicted and empirical fraction of time on platform across blocks of 3 trials for RW models assuming learning on shock trials (red), avoid trials (blue), or both trial types (purple; hybrid). B. Comparison of mean squared residuals per animal for each model type (One-way ANOVA, N=9). C. Observed behavior (red dots) and average predictions from hybrid Rescorla-Wagner model of PMA (black line) during blocks of 3 trials across both days of learning for shocked animals (N=9). D. Example schematic of trajectory distance (D1) and mean euclidean distance (D2) analyses of latent trajectories. Each trajectory spans 5s, starting at a colored circle and ending at an arrow marker. E. Example latent trajectories of average shock (red) and average avoid (blue) population responses and associated plots of correlation between mean Euclidean distance and PMA learning rate during tone onset epoch on day 1 (Pearson’s correlation, N=7). F. Same as E for pre-shock epoch on day 1. (Pearson’s correlation, N=7). G. Correlation plots of tone onset trajectory distance and PMA learning rates for shocked animals on day 1 aligned to trials of interest (Pearson’s correlation, N=8). H. Same as G for pre-shock epoch on day 1 (Pearson’s correlation, N=8). *P<0.05, **P<0.01. Circles represent female subjects and diamonds represent male subjects. Graphs show mean ± SEM. See Supplemental Table 1 for additional statistical information.

To understand how PL population dynamics relate to individual learning rates, we used a dimensionality reduction approach called calcium imaging linear dynamical system (CILDS, Koh et al., 2023). We focused on activity during two key behavioral epochs - the tone-onset epoch (0-5s) when the threat level increases, and the pre-shock epoch (23-28s) when shock is imminent, and decoding was most significant (Figure 3). For each epoch, we used CILDS to determine PL activity trajectories in latent space and then examined how aspects of PL dynamics (Figure 4D) related to individual learning rates.

Well trained mice have specific, tone-induced population activity patterns that are associated with avoidance actions (Ehret et al., 2024; Jercog et al., 2021). We therefore hypothesized that the emergence of PL activity states that distinguish between successful and unsuccessful avoidance trials may be a key signature of learning. To examine this, we measured the mean framewise Euclidean distance between population trajectories on successful and unsuccessful trials (Figure 4D_1_) and correlated Euclidean distance with the ‘avoid’ learning rates from our hybrid model (Figure 4E,F). We focused our analysis on day 1, when mice were rapidly learning but behavioral performance had not yet stabilized. The mean Euclidean distance during the tone onset epoch was significantly correlated with learning rate (Figure 4E). In contrast, we observed only trend-level, non-significant correlations during the pre-shock epoch (Figure 4F). These findings suggest that the rapid emergence of tone-induced representations of avoidance behaviors is a key signature of learning.

The data in Figure 4F average activity across trials and thus may miss effects limited to specific trials. Mice learn PMA rapidly and our modeling data suggest that mice learn both from failed and successful trials (Figure 4A-C). Thus, the first shock and first successful avoidance are likely highly salient drivers of learning that we predicted would be reflected in latent PL activity states. We therefore examined relationships between population dynamics and learning rates on individual trial types including the first shock trial, the first successful trial, and the subsequent trials, respectively. Of note, the first shock trial represents the moments just before the first foot shock, before the animals learned anything about aversive outcomes.

We measured population dynamics using the Euclidean distance between the start and end points of each trajectory (‘Trajectory Distance’; Figure 4D_2_). During the tone onset epoch, we found no significant correlations between trajectory distance and learning rate (Figure 4G). During the pre-shock epoch, trajectory distance had a strong positive correlation with learning rate on the first avoidance trial, but not on the first shock trial. Taken together, these results indicate that faster learning is associated with 1) with greater discrimination of trial outcome immediately after the tone onset and 2) with enhanced PL dynamics during the pre-shock epoch, when danger is high because the shock is imminent.

To better understand these changes in population dynamics, we compared shocked and non-shocked mice (Figure S4). We observed significant differences between shocked and NS control mice only on the trial following the first successful avoidance. During the tone-onset epoch, trajectory distance was significantly higher in shocked mice, whereas for the pre-shock epoch it was significantly lower than NS controls (Figure S4C,D). These differences were not present when we aligned population vectors to later trials (last shock; Figure S4). Moreover, we observed no behavioral differences between shocked and non-shocked mice during either the tone onset (Figure S4E) or during the pre-shock epoch (Figure S4F). This suggests the differences in population activity following the first successful trial reflect a learning process rather than differences in avoidance or freezing behaviors.

## DISCUSSION

In this study, we combined PMA with miniscope recordings to identify the transformations in PL activity that occur as animals learn to avoid signaled threats. At the single neuron level, we observed no changes in the proportion of significantly responsive PL neurons. However, by day 2 of PMA, shocked mice developed a greater proportion of neurons that exhibited mixed selectivity, encoding combinations of environmental cues and behaviors. Moreover, on day 2, the threatening tone strongly affected encoding of movements between safe and risky locations, and along the same lines, tone encoding was strongly impacted by the animal’s location.

Despite the lack of early changes in single neuron encoding, SVMs revealed that we could already decode trial outcomes from PL activity on Day 1 and that the predictive power was enhanced by using only the significantly responsive neurons. Finally, we uncovered specific changes in PL population activity that correspond to individual learning rates estimated using a Rescorla-Wagner model of PMA behavior. At the onset of the tone, the extent to which activity discriminated between trial outcomes was positively correlated with learning rates. During the pre-shock epoch, larger shifts in the population activity state were associated with faster learning rates, but these effects were transient and only apparent around the time of the first successful avoidance trial. Our detailed account of this sequence of events underlying rapid avoidance learning reveals how changes in the PL population code precede changes in the encoding properties of individual neurons, which emerge once the new behavioral strategy has stabilized.

### Changes in single cell encoding during rapid avoidance learning

Previous studies have described PL neurons that encode conditioned tones, foot shocks, and freezing behaviors during and after fear conditioning (Burgos-Robles et al., 2009; Courtin et al., 2014; Cummings et al., 2022; Cummings & Clem, 2020; Kitamura et al., 2017). A previous study of rats that had fully learned PMA reported a greater number of PL neurons that alter their firing (either increase or decrease) following tone onset and more PL cells that decrease their firing following avoidance actions compared to NS controls (Diehl et al., 2018). In line with this, we identified individual PL neurons that encoded the tone, platform entries, exits and location, and freezing behavior. We found that the overall proportion of significantly responsive neurons was similar in shocked vs. NS mice, indicating that changes in the number of behaviorally modulated PL neurons is not required for learning to avoid threats. Instead, we observed more neurons with mixed selectivity in mice that learned PMA compared to controls. We also showed that the activity of neurons encoding actions was highly modulated by threat level and that neurons encoding the tone were highly modulated by the animals’ location. We built on previous studies by revealing when learning-related changes first emerge. As these changes only emerge on day 2, they probably do not drive initial learning, but may stabilize learned associations to maintain learned behavioral strategies.

During threat assessment, mixed selectivity in PL has been proposed to reflect associative learning about a predictive cue combined with activation of appropriate behavioral strategies (e.g. active avoidance or defensive freezing) via the many downstream targets of PL (Grunfeld & Likhtik, 2018). Previous studies showed that during an avoidance task, mPFC has more mixed selective neurons compared to the basolateral amygdala (BLA). There is evidence that inputs from both BLA and the hippocampus influence mixed selectivity by sending information about conditioned stimuli and spatial cues, respectively (Burgos-Robles et al., 2017; Likhtik et al., 2014; Sotres-Bayon et al., 2012). These signals are likely processed within mPFC and then routed to appropriate top-down circuits for behavioral control (Gongwer et al., 2023; Kajs et al., 2022). For instance, activity in PL-BLA projections enhances threat avoidance whereas activity in PL projections to the nucleus accumbens (NAc) increases risky reward seeking during PMA (Diehl et al., 2020). Thus, PL-BLA neurons may contribute to the elevated activity during platform entries while PL-NAc neurons may contribute to the enhanced activity during platform exits on day 2, either promoting risky exploration during the tone or reporting the impending danger. While our study recorded broadly from PL neurons, future studies can examine this by performing miniscope recordings from anatomically defined neuronal classes during threat avoidance learning.

### Signatures of PL population activity that emerge early in learning

Neural population trajectories can reveal latent processes that are difficult to address by only analyzing single cell activity (Herry & Jercog, 2022; Sylte et al., 2024). PL population activity represents high level aspects of behavior such as strategic choices in a decision making task (Powell & Redish, 2014), integration of past decisions with future rewards (Herry & Jercog, 2022; Siniscalchi et al., 2019; Sylte et al., 2024), and threat-induced avoidance behaviors but not comparable movements in the absence of threatening cues (Ehret et al., 2024). But how and when new population codes emerge in PL following the events that drive aversive learning has not been well studied. Here we found that shifts in PL population dynamics occur earlier than changes in single neuron dynamics. Changes in the PL population code were evident following the first successful avoidance trial, before a stable pattern of avoidance behavior emerged.

We focused our population analyses on two key epochs during the tone: 1) the tone onset, which marks a rapid increase in threat level and 2) the epoch immediately preceding the footshock, when animals must engage in avoidance behavior to prevent encounters with the shock. Activity in each epoch has been shown to have different relationships with behavior. For instance, in well-trained animals, inhibiting mPFC activity during the tone onset delayed avoidance while inhibiting later in the tone decreased the probability of avoiding (Diehl et al., 2020; Jercog et al., 2021). Also, population activity immediately preceding the shock, but not at the onset of the conditioned tones, was an accurate predictor of trial outcome (Jercog et al., 2021). Our results using SVM decoders were consistent with these findings, showing that decoder performance is maximal during the pre-shock epoch. We also built on this literature by showing 1) that high decoder performance was already present on day 1, before animals established a stable avoidance strategy, and 2) that the significantly responsive neurons identified using ROC strongly contributed to the ability to decode trial outcomes on day 1. This suggests that during rapid learning, PL draws upon a pool of already tuned neurons to establish new patterns of population activity that help animals link threatening cues with avoidance behaviors.

By combining our Rescorla-Wagner model of behavior with CILDS dimensionality reduction, we determined how PL dynamics unfold across individual trials and identified the aspects of PL dynamics that reliably track with individual learning rates. Compared to principal component analysis, CILDS, which uses a linear dynamics systems model of Ca^2+^ activity, is uniquely suited for examining how PL dynamics evolve across trials. CILDS incorporates trial structure into the model and allows time series modeling, which is more sensitive for detecting changes across short time spans or in data that is not well represented by averages (Cunningham & Yu, 2014; Koh et al., 2023; Valente et al., 2022). Using this novel approach, we found that during the tone onset epoch, the extent to which activity patterns distinguished between trial outcomes was positively correlated with learning rate. During the pre-shock epoch, larger changes in PL population activity were associated with faster learning rates, but only on the trial with the first successful avoidance. Our data suggest that PL dynamics around the first successful trial may be particularly salient for learning. While our findings are correlative, there is a vast literature showing that PL activity is required for signaled threat avoidance (Bravo-Rivera et al., 2014; Capuzzo & Floresco, 2020; Diehl et al., 2018; Fernandez-Leon et al., 2021). Our studies build a foundation for future work that uses precisely timed manipulations on individual trials to investigate the causal roles of PL activity patterns in rapid avoidance learning.

## STAR METHODS

### Resource Availability

#### Lead contact

Further information and requests for raw data and reagents will be fulfilled by the lead contacts, Dr. Laura DeNardo & Dr. Scott Wilke.

#### Materials availability

This study did not generate new unique reagents.

#### Data and code availability

- All data reported in this paper will be shared by the lead contact upon request.
- All original code has been deposited at Zenodo and is publicly available as of the date of publication. DOIs are listed in the key resources table.
- Any additional information required to reanalyze the data reported in this paper is available from the lead contact upon request

### Experimental Model and Subject Details

#### Animals

Female and male C57B1/6J mice (JAX Stock No. 000664) aged 10–16 weeks were group housed (2–5 per cage) and kept on a 12 hr light cycle. 6 of 11 shocked mice were female while 5 of 7 non-shocked mice were female. After GRIN lens implantation, all animals were single housed until the end of behavioral testing. Following the baseplate surgery, all animals were handled daily and habituated to the weight and feel of wearing the miniscope for 10 days. All animal procedures followed animal care guidelines approved by the University of California, Los Angeles Chancellor’s Animal Research Committee.

### Method Details

#### Behavior video recordings

Behavioral videos were acquired at 30fps using a ELP 2.8–12 mm Lens Varifocal Mini Box 1.3 megapixel USB Camera.

#### Platform-mediated avoidance

For all PMA sessions, a conditioning chamber was used consisting of an 18cm x 18cm x 30 cm cage with a grid floor wired to a scrambled shock generator (Lafayette Instruments) surrounded by a custom-built acoustic chamber. The chamber was scented with 50% Windex. One corner was covered with a thin acrylic platform (3.5in x 4in x 0.5in), amounting to 25% of the chamber floor. During the first day of PMA, mice were presented with three baseline 30s 4 kHz tones (CS), followed by nine presentations of the CS that co-terminated with a 2s footshock (0.14mA). The following day, mice were presented with twelve CS that coterminated with a shock.

#### Viruses

AAV1-syn-jGCaMP7f.WPRE (ItemID: 104488-AAV1) was purchased from Addgene and diluted to a working titer of 8.5×1012 GC/ml.

#### Miniscope surgery and baseplating

For miniscope recordings, all mice underwent two stereotaxic surgeries (Cai et al., 2016). First, adult WT mice were anesthetized with isoflurane and secured to a stereotaxic frame (Kopf 963). Mice were placed on a heating blanket and artificial tears kept their eyes moist throughout the surgery. After exposing the skull, a burr hole was drilled above PL in the left hemisphere (+1.85, –0.4, –2.1 mm from bregma). A Hamilton syringe containing AAV1-Syn-jGCaMP7f-WPRE was lowered into the burr hole and 600 nL of AAV was pressure injected using a microinjector (WPI, UMP3T-1). The syringe was left in place for 10 min to ensure the AAV did not spill out of the target region and then the skin was sutured. After recovery, animals were housed in a regular 12 hr light/dark cycle with food and water ad libitum. Carprofen (5 mg/kg) was administered both during surgery and for 2 days after surgery together with amoxicillin (0.25 mg/mL) for 7 days after surgery. One week later, mice underwent a GRIN lens implantation surgery. After anesthetizing the animals with isoflurane (1–3%) and securing them to the stereotaxic frame, a 1mm craniotomy was made above the virus site, and the cortical tissue above the targeted implant site was carefully aspirated using 27-gauge and 30-gauge blunt needles. Buffered ACSF was constantly applied throughout the aspiration to prevent tissue desiccation. The aspiration ceased after full termination of bleeding, at which point a GRIN lens (1 mm diameter, 4 mm length, Inscopix 1050–002176) was stereotaxically lowered to the targeted implant site (– 2.0 mm dorsoventral from skull surface relative to bregma). Cyanoacrylate glue was used to affix the lens to the skull. Then, dental cement sealed and covered the exposed skull, and Kwik-Sil covered the exposed GRIN lens. Carprofen (5 mg/kg) and dexamethasone (0.2 mg/kg) were administered during surgery and for 7 days after surgery together with amoxicillin (0.25 mg/mL) in the drinking water. Two weeks after implantation, animals were anesthetized again with isoflurane (1–3%), and a miniscope attached to an aluminum baseplate was placed on top of the GRIN lens. After searching the field of view for in-focus cells, the baseplate was cemented into place, and the miniscope was detached from the baseplate. A plastic cap was locked into the baseplate to protect the implant from debris and allow the baseplate to set.

#### Miniscope recordings

Mice were handled and habituated to the weight of the microscope for 10 days before behavioral acquisition. On the recording day, a V4 miniscope was secured to the baseplate with a set screw and the mice were allowed to acclimate in their home cage for 5 min. Imaging through the miniscope took place throughout the entire PMA session (∼30 min) and retrieval (∼18 min) sessions on following days. Behavior was simultaneously recorded using miniscope recording software to synchronize the data streams (https://github.com/Aharoni-Lab/Miniscope-DAQ-QT-Software).

#### Miniscope data processing

Frames in which animals were freezing and/or on the safety platform were determined using BehaviorDEPOT. Cell footprints and Ca2+ fluorescence time series were extracted from miniscope recordings using Minian (https://github.com/denisecailab/minian). Videos were recorded at 30Hz and were spatially and temporally downsampled by by a factor of 2. Accepted neurons and their calcium activity traces were exported to MATLAB for further analysis using custom scripts. We identified 1394 neurons across 9 mice in Shock animals and 847 neurons across 7 mice in Non-shock animals. Custom MATLAB software was used to align data from the behavior camera and the miniscope camera.

#### ROC Analysis

To identify neurons that were active during features relevant to PMA, we evaluated single cell responses to the conditioned tones, freezing behavior, and the safety platform. We defined behaviorally-modulated cells as those that had a significant modulated response to any of the following: tone onsets or offsets (first or last 5s of tone), bouts of freezing behavior, or platform entries or exits (from 1s before entry/exit until 4s after). We plotted receiver operating characteristic curves (ROC) for individual neurons and measured the area under the curve (auROC). ROCs plot the true positive rate (true positive/(true positive + false negative)) against the false positive rate (false positive/(false positive + true negative)) over a range of probability thresholds. Neurons with high auROC values therefore predict the behavioral variable of interest with a high true positive rate and low false positive rates over a large range of thresholds. To determine if a neuron significantly encoded a particular behavioral event, we generated a null distribution of auROCs by circularly shuffling event timing and recalculating the auROC over 1000 permutations. Neurons were considered significantly activated during a behavior if their auROC was greater than 97.5% of auROCs in the null distributions and significantly suppressed if their auROC value was in the lowest 2.5% of all auROCs in the null distribution.

*Cells Analyzed for* Figure 1 *Heatmaps:*

Excited:

– Day 1: (ToneOn: Shk, N=61; NS, N=24; Frz: Shk, N=162; NS, N=137; PlatEnt: Shk, N=150; NS, N=95; PlatExt: Shk, N=131; NS, N=66).

– Day 2: (ToneOn: Shk, N=34; NS, N=34; Frz: Shk, N=171; NS, N=150; PlatEnt: Shk, N=112; NS, N=102; PlatExt: Shk, N=104; NS, N=100).

Inhibited:

– Day 1: (ToneOn: Shk, N=52; NS, N=25; Frz: Shk, N=334; NS, N=187; PlatEnt: Shk, N=87; NS, N=66; PlatExt: Shk, N=78; NS, N=67).

– Day 2: (ToneOn: Shk, N=37; NS, N=22; Frz: Shk, N=283; NS, N=198; PlatEnt: Shk, N=80; NS, N=92; PlatExt: Shk, N=74; NS, N=82).

### Frequency Distributions of auROC Values

To identify differences in response modulation to intersections of tone and platform activity per animal, we collected auROC values for all cells responding to a particular feature and separated them by learning event (tone on/off; on/off platform). Animals were excluded if they had fewer than 3 ROC-identified cells in a particular event category. All auROC values were collected, and inhibited values (auROC = 0:0.5) were converted into a fixed range using the formula: 1 - auROC. These auROC values were then used to construct cumulative frequency histograms for each analysis. To measure modulation strength, we derived a modulation score from the AUC of each animal’s frequency histogram using the formula: Modulation Score = 1 - AUC.

### Decoding Analysis

To determine how well the PL activity predicts avoidance in response to shock, we employed a support vector machine (SVM) with a radial basis function (RBF) kernel (Chang & Lin, 2011) to decode trial outcomes from population activity. We determined the outcome to decode from each trial as either a successful avoidance (1) or a failed shock (0). We excluded the first three trials from the baseline period of day 1 as PMA learning had not yet begun. We used the ΔF/F_o_ activity of the units in each trial as features to train and test the decoder. The activity of single units in each trial was binned in 200ms time windows. To evaluate how well PL activity predicts avoidance behavior before, during, and after the shock, we trained an SVM on the activity of each animal and tested it across day 1. SVMs for each animal were trained on binned activity using ‘leave-one-out’ stratified cross-validation. To maximize the decoder performance, we optimized the RBF SVM regularization parameters for each time bin and each animal, specifically the misclassification cost parameter, *C*, and the data complexity parameter, *γ,* by using a grid search with five fold cross-validation. The SVM performance for each bin was calculated as the min-max normalized F1 score, averaged across subjects. The F1 score was calculated as follows: F1 = 2* [precision * recall / (precision + recall)], where precision = True positives / (False positives + True positives) and recall = True positives / (False negatives + True positives).

We averaged the SVM performance over four epochs per mouse to compare the decoding performance between epochs and between groups (Shock, Non-shock): “Cue-onset” (0 - 5 s), “Pre-shock” (23 - 28 s), “Shock” (28 - 33 s), and “Post-shock” (34 - 38 s).

### Dimensionality Reduction of Recorded Neural Data

We used a dimensionality reduction pipeline for calcium imaging data, CILDS (https://github.com/kohth/cilds), to extract 3 latent variables from each session that were trained using neural calcium data during 5s windows of the tone (Tone onset: 1 to 5s; Pre-shock: 23 to 28s) for each trial of a session. Extracted latent variables were averaged by trial outcome or aligned to events of interest. Two sessions were excluded due to technical issues running the algorithm on those datasets.

Distance was assessed using 3 measures:

– *Trajectory distance*: the Euclidean distance between the trajectory start and end points
– *Mean Euclidean distance*: average value of framewise Euclidean distances calculated between each point in a set of compared trajectories for each animal

### Rescorla-Wagner Modeling

To correlate neural dynamics during PMA to mechanistic changes in subjects’ behavior across learning, a Rescorla-Wagner learning model was fit to the proportion of trial time spent on the platform for each mouse across trials. The proportion of total trial time spent on the safety platform during conditioned tones was modeled using a Rescorla-Wagner model (Rescorla & Wagner, 1972). We seeded the model with the proportion amount of baseline time animals spent on the platform prior to shock. Because mice in this task experience a mixture of punishment (shock trials) and negative reinforcement (successful escapes to the platform), we used a version of model with two complimentary learning rates, α_failure_, which weights the rate of change in the platform value following an unsuccessful trial, and α_success_, which weights the rate of change in the platform value following a successful avoid. It is assumed that the subjects’ behavioral expression of platform value is the proportion of trial time spent on the platform.

Models were fit to individual subject data, concatenated across Days 1 and 2, with custom Python code using maximum likelihood estimation. The trial-by-trial change in the value of the platform was calculated as:

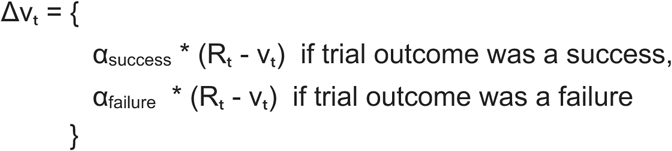

Where:

– α_success_ = 1 - α_failure_
– Δvₜ is the change in associative strength of the platform at trial t
– α is the learning rate (free parameter)
– Rₜ is the outcome at trial t
– vₜ is associative strength at time t (initially the baseline probability of being on platform)

### Histology

Mice were transcardially perfused with phosphate-buffered saline (PBS) followed by 4% paraformaldehyde (PFA) in PBS. Brains were dissected, post-fixed in 4% PFA for 12–24 hr and placed in 30% sucrose for 24–48 hr. They were then embedded in Optimum Cutting Temperature (OCT, Tissue Tek) and stored at –80 °C until sectioning. 60 µm floating sections were collected into PBS. Sections were washed 3×10 min in PBS and then blocked in 0.3% PBST containing 10% normal donkey serum (Jackson Immunoresearch, 17-000-121) for 2 hr. Sections were then stained with chicken anti-GFP primary antibody (Aves 1020, 1:2000) in 0.3% PBST containing 3% donkey serum overnight at 4 °C. The following day, sections were washed 3×5 min in PBS and then stained with secondary antibody (goat anti-chicken 488, 1:500) in 0.3% PBST containing 5% donkey serum for 2 hr at room temperature. Sections were then washed 5 min with PBS, 15 min with PBS + DAPI (Thermofisher Scientific, D1306, 1:4000), and then 5 min with PBS. Sections were mounted on glass slides using FluoroMount-G (ThermoFisher, 00-4958-02) and then imaged at 10x with a Leica slide scanning microscope (VT1200S).

### Quantification and Statistical Analysis

Data analyses were performed using custom scripts written in MATLAB, Python, and Prism. All statistical analyses were performed in GraphPad Prism. Behavior data (Figure 1) and decoding data (Figure 3) were analyzed with a two-way repeated measures ANOVA with Geisser-Greenhouse correction and post hoc Šidák’s multiple comparisons test. Cumulative frequency histograms (Figure 2) were analyzed by ordinary two-way ANOVA and Fisher’s LSD. RW model squared residuals (Figure 4) were analyzed with a repeated measures one-way ANOVA with Geisser-Greenhouse correction. Chi squares were used to compare pie charts in Figure 1. All other comparisons were made using a t-test with Welch’s correction. Further statistical information can be found in Supplemental Table 1.

### Data and Software Availability

Analyzed datasets are available on Zenodo (10.5281/zenodo.13381210). Custom data analysis scripts were coded in MATLAB or Python and are available on Zenodo (10.5281/zenodo.13351302).

## Acknowledgments

We thank Dean Buonomano for critical feedback and Daniel Aharoni and Federico Sangiuliano for technical support building and troubleshooting miniscopes. This work was supported by R01MH137461 (L.A.D, S.A.W), R01MH127214 (L.A.D.), R01MH131858 (S.A.W.), F32MH133387 (Z.Z.), T32NS115753 (T.G.), and an Achievement Rewards for College Scientists Predoctoral Fellowship (C.J.G).

## Author Contributions

Conceptualization, C.J.G., S.A.W., & L.A.D.; Methodology, C.J.G., T.G., & A.S.; Software, C.J.G., T.G., & A.S.; Formal Analysis, C.J.G., T.G., & A.S.; Writing – Original Draft, C.J.G. & L.A.D.; Writing – Review & Editing, C.J.G., T.G., A.S., Z.Z., S.A.W., & L.A.D.; Supervision, S.A.W. & L.A.D; Funding Acquisition, S.A.W. & L.A.D.

