## Supplemental Material for "Transformations in prefrontal ensemble activity underlying rapid threat avoidance learning"

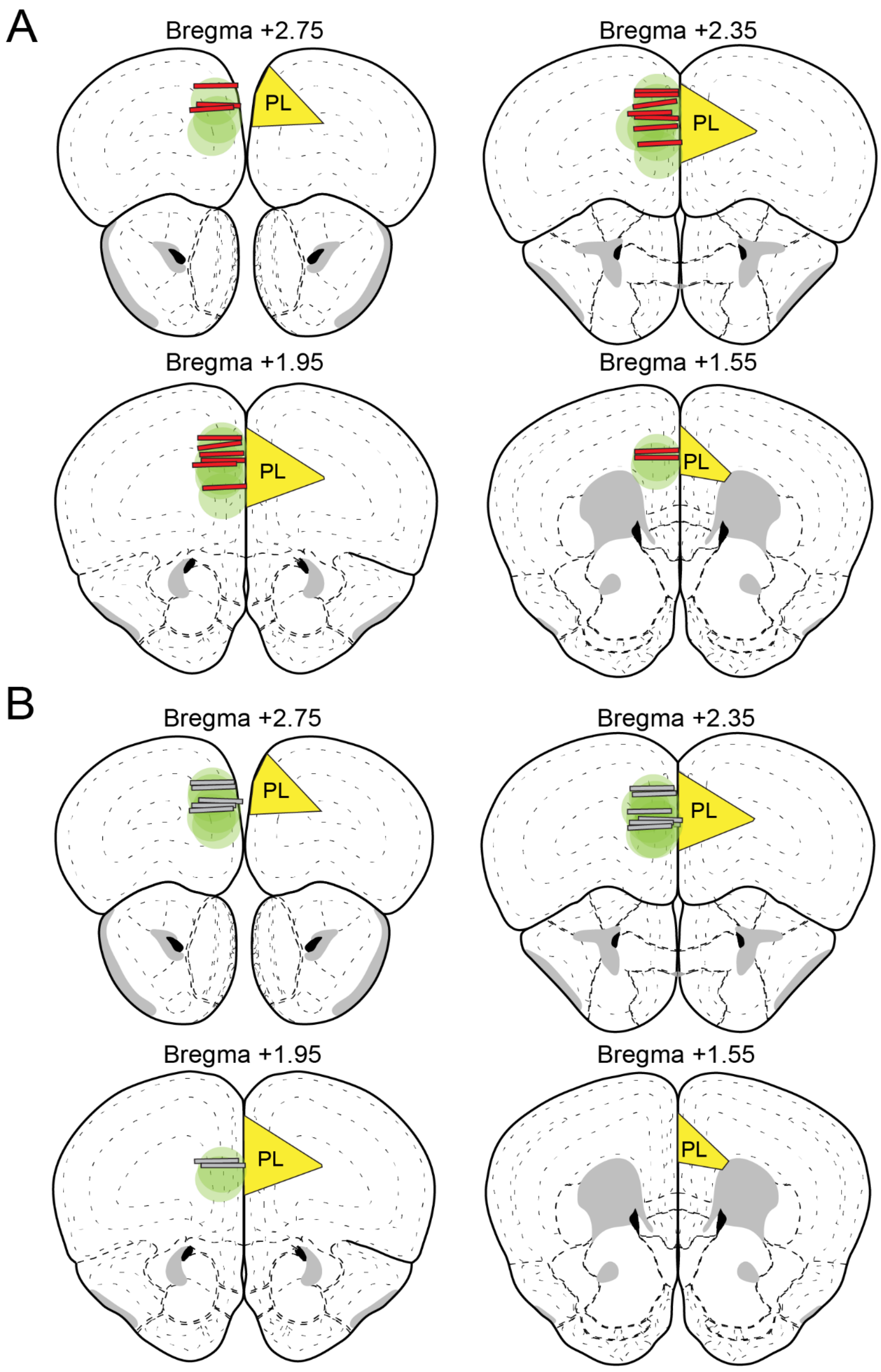


**Figure S1. GRIN Lens Placement and GCaMP Expression Sites**

A. Coronal sections of mPFC showing the placement of GRIN lens edges and GCaMP expression for animals in the shock group. Red bars show location of lens edge and green area shows GCaMP expression region.

B. Same as A for non-shocked animals.

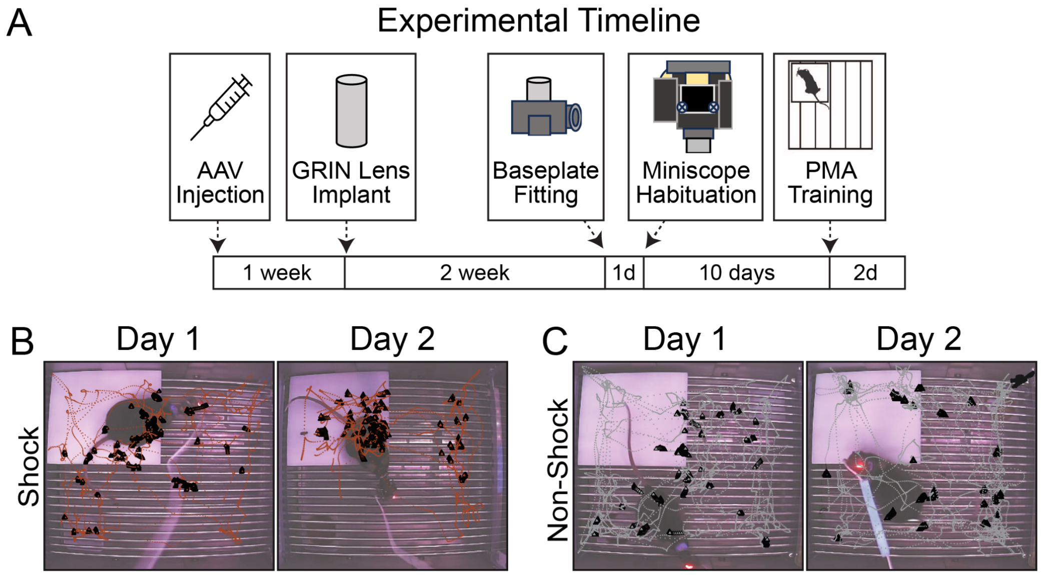


**Figure S2. Details of Experimental Design**

A. Experimental timeline for miniscope surgeries and platform-mediated avoidance.

B. Example maps of shocked animal trajectory during PMA days 1 *(left)* and 2 *(right)*. Plots show each animal's location during tone periods for all trials across each session. Black triangles show the location of animal freezing.

C. Same as B for a non-shocked mouse.

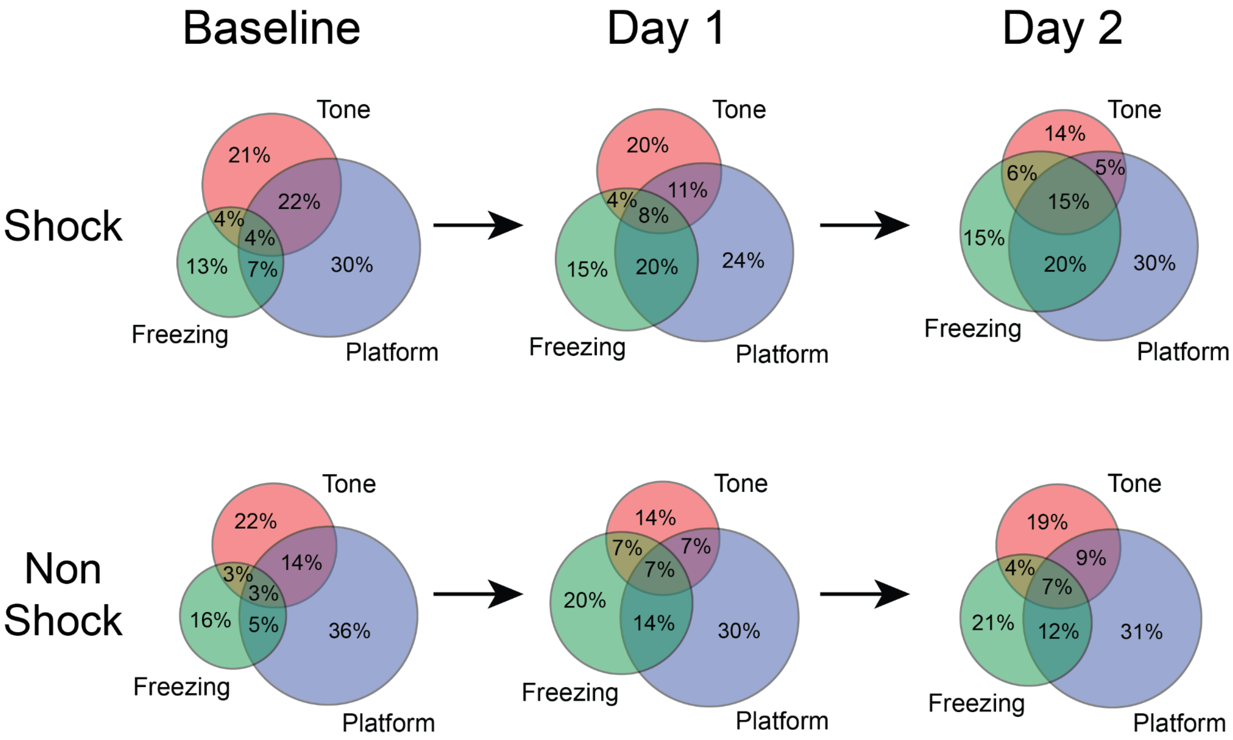


**Figure S3. Mixed Encoding of PMA Features Across Learning**
Venn diagrams showing breakdown of responsive cells during baseline, day 1 (last 3 tones), and day 2 (last 3 tones). Values show percentage of feature responsive cells identified at each period (as indicated in Figure 1K-M). Scaling is approximate.

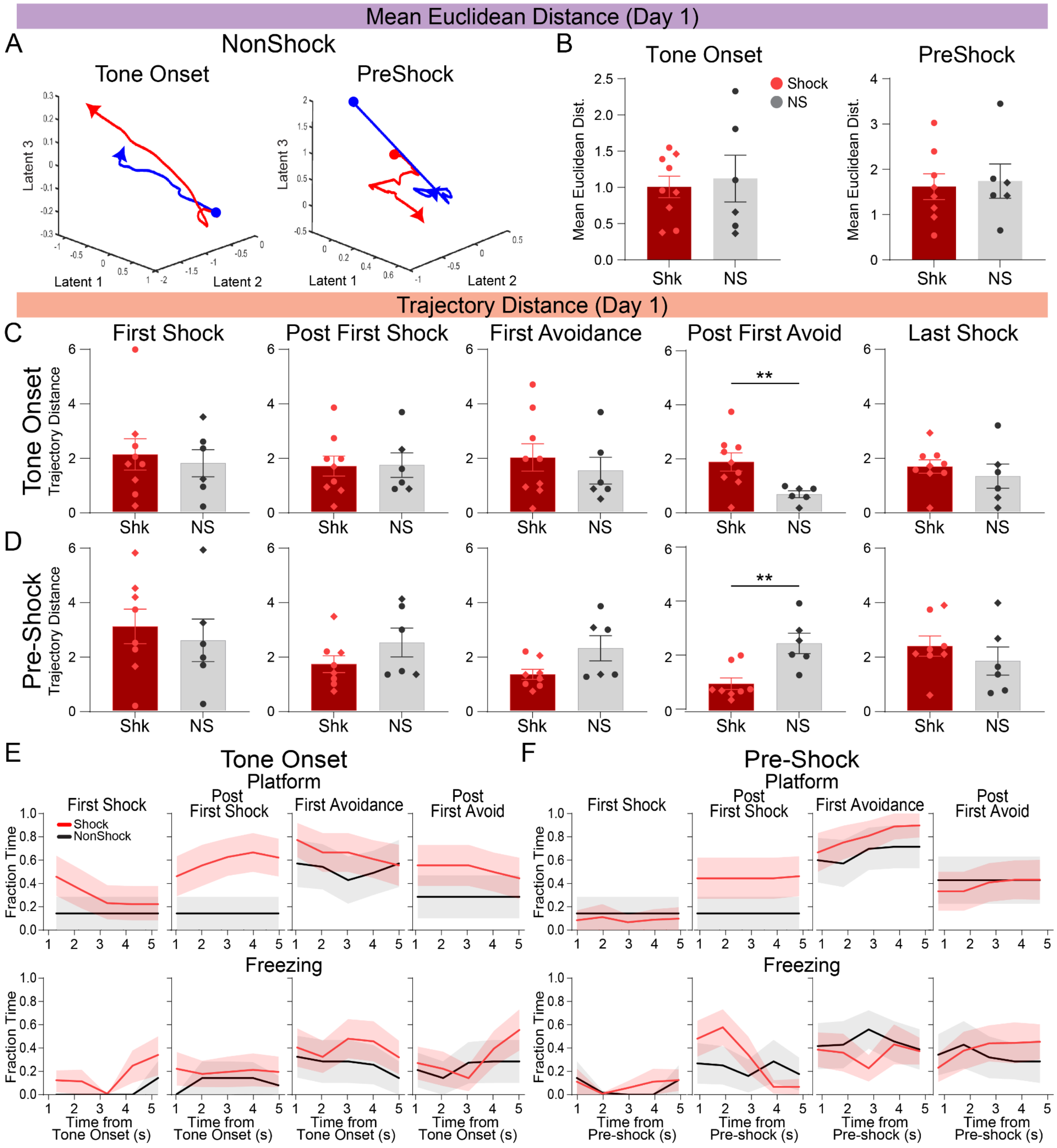


**Figure S4. Comparing PL Population Dynamics in Shocked and Non-shocked Mice**

A. Example 3D trajectories of average shock (red) and average avoid (blue) population responses during tone onset and pre-shock epochs from representative non-shocked example mice on day 1.

B. Mean Euclidean distance between average shock and avoid trials for tone onset and pre-shock epochs. (Welch's t-test; ToneOn: Shk, N=9; NS, N=6; Pre-Shock: Shk, N=8; NS, N=6).

C. Day 1 tone onset trajectory distance on individual trials (Welch's t-test; Shock, N=9; NS, N=6).

D. Same as D for pre-shock epoch (Welch's t-test; Shock, N=8; NS, N=6).

E. Mean fraction time on platform *(top)* and freezing *(bottom)* during the tone onset for shocked and non-shocked mice aligned to trials of interest on day 1 (Two-way ANOVA; Shock: N=9, NS: N=7).

F. Same as E for pre-shock epoch. (Two-way ANOVA; Shock: N=9, NS: N=7).

*P<0.05, **P<0.01. Graphs show mean + SEM. See Supplemental Table 2 for additional statistical information.

**Supplemental Table 1. Expanded ANOVA Results.**

Two-way repeated measures ANOVA with Geisser-Greenhouse correction and post hoc Šídák's multiple comparisons test

| **Figure 1** | Day | $F_{trial}$ | DFn, DFd | p-value | $F_{group}$ | DFn, DFd | p-value | $F_{interaction}$ | DFn, DFd | p-value |
|---|---|---|---|---|---|---|---|---|---|---|
| 1E: Time on Platform | 1 | 14.07 | 2.473, 44.51 | <0.0001 | 2.429 | 1, 18 | 0.1365 | 1.089 | 3, 54 | 0.3617 |
| | 2 | 0.654 | 2.375, 35.62 | 0.5514 | 35.87 | 1, 15 | <0.0001 | 1.970 | 3, 45 | 0.1320 |
| 1F: Successful Trials | 1 | 8.814 | 1.883, 33.90 | 0.0010 | 7.372 | 1, 18 | 0.0142 | 2.173 | 3, 54 | 0.1019 |
| | 2 | 0.7388 | 2.311, 34.67 | 0.5034 | 29.41 | 1, 15 | <0.0001 | 1.798 | 3, 45 | 0.1611 |
| 1G: Overall Freezing | 1 | 15.02 | 2.346, 42.22 | <0.0001 | 0.02244 | 1, 18 | 0.8826 | 1.960 | 3, 54 | 0.1309 |
| | 2 | 0.5929 | 2.174, 32.60 | 0.5720 | 13.98 | 1, 15 | 0.0020 | 1.000 | 3, 45 | 0.4014 |
| 1H: Latency | 1 | 3.567 | 2.674, 48.12 | 0.0247 | 0.7463 | 1, 18 | 0.399 | 1.320 | 3, 54 | 0.2773 |
| | 2 | 0.2622 | 2.574, 38.61 | 0.8230 | 19.25 | 1, 15 | 0.0005 | 2.503 | 3, 45 | 0.0713 |

Ordinary two-way ANOVA and Fishers LSD

| **Figure 2** | Day | $F_{tone/platform}$ | DFn, DFd | p-value | $F_{group}$ | DFn, DFd | p-value | $F_{interaction}$ | DFn, DFd | p-value |
|---|---|---|---|---|---|---|---|---|---|---|
| 2D: Platform Entries | 1 | 45.23 | 1, 24 | <0.0001 | 1.534 | 1, 24 | 0.2274 | 2.339 | 1, 24 | 0.1392 |
| | 2 | 35.87 | 1, 22 | <0.0001 | 5.355 | 1, 22 | 0.0304 | 0.7200 | 1, 22 | 0.4053 |
| 2F: Platform Exits | 1 | 78.37 | 1, 24 | <0.0001 | 0.05099 | 1, 24 | 0.8233 | 0.2518 | 1, 24 | 0.6204 |
| | 2 | 72.03 | 1, 20 | <0.0001 | 10.27 | 1, 20 | 0.0044 | 8.404 | 1, 20 | 0.0089 |
| 2J: Tone Onset | 1 | 0.1350 | 1, 24 | 0.7165 | 1.538 | 1, 24 | 0.2269 | 1.566 | 1, 24 | 0.2228 |
| | 2 | 0.9529 | 1, 18 | 0.3419 | 0.08758 | 1, 18 | 0.7707 | 6.751 | 1, 18 | 0.0182 |
| 2L: Tone Offset | 1 | 0.08148 | 1, 25 | 0.7777 | 0.2890 | 1, 25 | 0.5956 | 12.93 | 1, 25 | 0.0014 |
| | 2 | 6.513 | 1, 13 | 0.0241 | 0.6236 | 1, 13 | 0.4439 | 13.98 | 1, 13 | 0.0025 |

Two-way repeated measures ANOVA with Geisser-Greenhouse correction and post hoc Šídák's multiple comparisons test

| **Figure 3** | Day | $F_{period}$ | DFn, DFd | p-value | $F_{group}$ | DFn, DFd | p-value | $F_{interaction}$ | DFn, DFd | p-value |
|---|---|---|---|---|---|---|---|---|---|---|
| 3A: All Cells | 1 | 1.302 | 2.313, 27.76 | 0.2908 | 16.99 | 1, 12 | 0.0014 | 1.375 | 3, 36 | 0.2660 |
| 3B: BM Cells | 1 | 2.179 | 2.278, 27.33 | 0.1267 | 20.46 | 1, 12 | 0.0007 | 4.440 | 3, 36 | 0.0094 |
| 3C: Non Modulated Cells | 1 | 0.9164 | 1.913, 22.95 | 0.4101 | 5.988 | 1, 12 | 0.0308 | 0.6529 | 3, 36 | 0.5864 |

Repeated measures one-way ANOVA with Geisser-Greenhouse's epsilon

| **Figure 4** | Day | $F_{model}$ | DFn, DFd | p-value | $F_{subject}$ | DFn, DFd | p-value |
|---|---|---|---|---|---|---|---|
| 4B: Model Residuals | | 9.166 | 1.347, 10.78 | 0.0081 | 1.863 | 8, 16 | 0.1377 |

### Supplemental Table 1. Expanded ANOVA Results

**Supplemental Table 2. Expanded Behavior ANOVA Results.**

Two-way repeated measures ANOVA with Geisser-Greenhouse correction and post hoc Šídák's multiple comparisons test

| Figure 4S | Type | $F_{time}$ | DFn, DFd | p-value | $F_{group}$ | DFn, DFd | p-value | $F_{interaction}$ | DFn, DFd | p-value |
|---|---|---|---|---|---|---|---|---|---|---|
| Tone Onset (Platform) | First Shock | 1.558 | 1.458, 20.41 | 0.2333 | 0.5571 | 1, 14 | 0.4678 | 1.558 | 4, 56 | 0.1981 |
| | Post First Shock | 0.9601 | 1.633, 22.87 | 0.3812 | 4.213 | 1, 14 | 0.0593 | 0.9601 | 4, 56 | 0.4366 |
| | First Avoidance | 1.259 | 2.332, 32.65 | 0.3010 | 0.3062 | 1, 14 | 0.5887 | 1.089 | 4, 56 | 0.3710 |
| | Post First Avoid | 0.7656 | 1.000, 14.00 | 0.3963 | 0.8871 | 1, 14 | 0.3622 | 0.7656 | 4, 56 | 0.5521 |
| Tone Onset (Freezing) | First Shock | 1.807 | 1.574, 22.03 | 0.1918 | 2.878 | 1, 14 | 0.1119 | 0.4542 | 4, 56 | 0.7689 |
| | Post First Shock | 0.1073 | 2.174, 30.44 | 0.9125 | 0.7510 | 1, 14 | 0.4008 | 0.2105 | 4, 56 | 0.9315 |
| | First Avoidance | 1.026 | 2.983, 41.76 | 0.3906 | 0.4397 | 1, 14 | 0.5190 | 0.3734 | 4, 56 | 0.8266 |
| | Post First Avoid | 1.974 | 1.412, 19.77 | 0.1732 | 0.1774 | 1, 14 | 0.6800 | 1.025 | 4, 56 | 0.4023 |
| Pre-Shock (Platform) | First Shock | 0.03586 | 1.042, 14.58 | 0.8615 | 0.1386 | 1, 14 | 0.7152 | 0.03586 | 4, 56 | 0.9975 |
| | Post First Shock | 0.7656 | 1.000, 14.00 | 0.3963 | 1.682 | 1, 14 | 0.2156 | 0.7657 | 4, 56 | 0.5521 |
| | First Avoidance | 2.275 | 1.402, 19.62 | 0.1409 | 0.4994 | 1, 14 | 0.4914 | 0.2406 | 4, 56 | 0.9141 |
| | Post First Avoid | 0.5021 | 1.496, 20.95 | 0.5595 | 0.02651 | 1, 14 | 0.8730 | 0.5021 | 4, 56 | 0.7343 |
| Pre-Shock (Freezing) | First Shock | 1.028 | 1.906, 26.68 | 0.3683 | 0.1051 | 1, 14 | 0.7506 | 0.3036 | 4, 56 | 0.8743 |
| | Post First Shock | 3.876 | 3.070, 42.98 | 0.0148 | 0.2193 | 1, 14 | 0.6468 | 3.301 | 4, 56 | 0.0169 |
| | First Avoidance | 0.1161 | 2.713, 37.98 | 0.9380 | 0.2477 | 1, 14 | 0.6264 | 0.8993 | 4, 56 | 0.4706 |
| | Post First Avoid | 0.5009 | 2.330, 32.62 | 0.6385 | 0.07471 | 1, 14 | 0.7886 | 1.083 | 4, 56 | 0.3738 |

### Supplemental Table 2. Expanded Behavior ANOVA Results
